# Ocean acidification alters hypoxia sensitivity and oxyregulation in reef-building corals

**DOI:** 10.64898/2026.04.22.718605

**Authors:** Rene M van der Zande, Kelly W Johnson, Sophie Littke, Verena Schoepf

## Abstract

Coastal marine ecosystems are increasingly threatened by multiple stressors such as ocean acidification and deoxygenation, but how these co-occurring stressors interact is often poorly understood. This is especially true for tropical coral reefs where deoxygenation is an emerging yet understudied threat. Using hypoxia response curves combined with rigorous pH control, we show that acidification alters hypoxia sensitivity and oxyregulation of reef-building corals in a species-specific manner: three species exhibited increased sensitivity to various degrees, while the fourth showed enhanced tolerance. Consequently, acidification pushes critical hypoxia thresholds into oxygen regimes already prevalent on reefs today, potentially driving shifts in community composition and accelerating risks to reef resilience as these stressors intensify in the future. Our findings challenge assumptions of uniform coral vulnerability under multi-faceted climate change, emphasizing the need for trait-based approaches and to account for stressor interactions in predictive models to better anticipate coral reef futures under rapid climate change.

## Introduction

Ocean deoxygenation and the emergence of dead zones are becoming increasingly prevalent and damaging to coastal marine ecosystems globally ^1–3^. Its effects likely already surpass those of ocean warming and acidification ^4^, prompting appeals from researchers to include aquatic deoxygenation in the Planetary Boundary Framework ^5,6^. Coral reefs, too, have experienced declines in seawater dissolved oxygen (DO) and the formation of local dead zones, resulting in widespread mortality and biodiversity loss ^7–9^. When deoxygenation co-occurs and interacts with other climate change stressors, ^10–12^, it can lead to devastating compound events that may be much more damaging than each stressor individually ^13^. However, the impacts of multiple stressors and compound events are poorly understood in many marine taxa, underscoring the need for more comprehensive research.

Although the effects of combined warming and deoxygenation on marine ecosystems have recently received considerable attention ^1,11,14^, the combined impacts of acidification and deoxygenation remain critically understudied, despite strong evidence of their detrimental effects on marine taxa ^10,15^. This is particularly true for tropical coral reefs where deoxygenation has long been overlooked as an emerging threat ^3,16,17^. Notably, corals (and other key marine taxa) were therefore poorly or not at all represented in these analyses ^1,10,15^, as the impacts of hypoxia on corals have only recently begun to be investigated. To date, no studies have investigated the effects of hypoxia and acidification beyond natural diel cycles of DO and pH in coral tissues (i.e., >12 hrs exposure), despite the fact that deoxygenation is accompanied by decreases in pH on both global and local reef scales ^8,10^. Thus, it remains currently unknown whether acidification alters coral hypoxia tolerance – an organism’s capacity to both regulate and maintain physiological functioning under decreasing oxygen saturation 10,15.

Crucially, the experimental approach that is widely used to determine an organism’s hypoxia threshold may be inherently confounded by acidification if low pH alters hypoxia sensitivity ^10,18^. Hypoxia tolerance thresholds are typically assessed using hypoxia response curves (Fig. 1) where organisms are incubated in sealed respirometry chambers without pH control and draw down DO over many hours until reaching anoxia. Due to the stoichiometric coupling of O2 and CO2 in metabolism, this automatically results in CO_2_ build-up and seawater acidification. Our data indicate that pH decreased by up to 0.7 units during the long incubations to reach anoxia (Supplementary Methods), a change driven by respiratory CO2 release rather than calcification during the incubation period. Yet, the potentially confounding effects of low pH on metabolically-derived hypoxia thresholds are at present unknown in corals as well as many other taxa ^10,18^.

**Figure 1.**
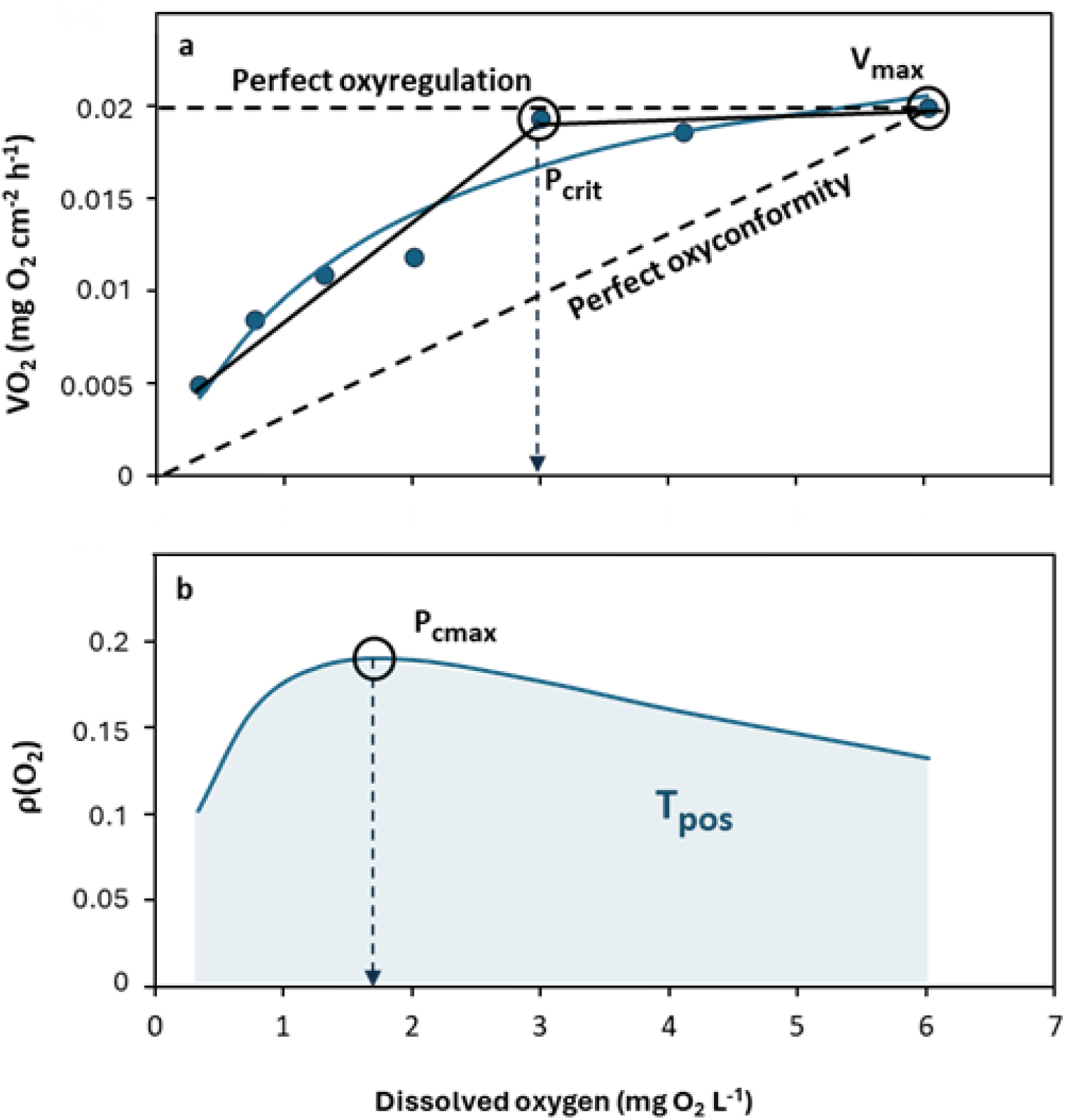
Schematic representation of a hypoxia response curve, its associated oxygen regulation profile, and different metrics of hypoxia sensitivity and regulation for one genotype of *Orbicella annularis* from this study. **(a)** Example of respiration rates (VO_2_) measured across the dissolved oxygen (DO) range from 0 to 100% air saturation (0 – 6.1 mg O_2_ L^-1^ at 30°C). Horizontal and diagonal black dashed lines represent perfect oxygen regulators and conformers, respectively. Maximum respiration rates (V_max_) are reached near maximum DO. The critical hypoxia threshold P_crit_ (dotted vertical black line; defined as the DO threshold at which VO_2_ can no longer be maintained constant) was obtained from segmented regression analysis and the calculation of the inflection point (solid black line). A fitted Michaelis-Menten function (solid teal line) was used to determine the relationship between DO and VO_2_ and to calculate V_max_, and to serve as the base for the establishment of the oxygen regulation function ρ(O_2_). **(b)** Maximum values for the regulation function ρ(O_2_) (solid teal line; same genotype) indicate the oxygen concentration where the maximum positive oxyregulation is applied to maintain metabolism (P_cmax_; dashed black line). The integral of the ρ(O_2_) function (shaded area) represents the total amount of positive oxyregulation (T_pos_). Note that the DO threshold values for P_crit_ (a) and P_cmax_ (b) represent different aspects of an organism’s response to low oxygen and do not necessarily coincide, as illustrated in this example.

Several key metrics to determine hypoxia sensitivity thresholds and (oxy)regulation of metabolism can be extracted from the fitted hypoxia response curves (Fig. 1) ^19,20^. The most widely used metric to determine hypoxia sensitivity is P_crit_, the critical oxygen threshold below which hypoxia prevents the maintenance of stable respiration rates (Fig. 1a). Oxyregulatory capacity can be characterized by the oxygen concentration where maximum positive regulation of metabolism is achieved (P_cmax_), and the maximum positive oxyregulation capacity (T_pos_) (Fig. 1b). To date, only few studies have characterized these thresholds in corals, revealing substantial variation in critical hypoxia thresholds and oxyregulation capacity between species and regions ^21–23^. However, most of these estimates derive from the hours-long, continuous drawdown incubations previously described ^21,22,24,25^, raising the possibility that our current knowledge of coral hypoxia tolerance is confounded by synergistic effects of hypoxia and acidification.

To address these knowledge gaps, we conducted hypoxia response curves with rigorous pH control to test whether acidification alters hypoxia sensitivity in four ecologically important coral species. Because coral responses to both hypoxia and acidification vary widely among species and genotypes ^26,27^, any interactive effects were expected to be highly species-specific, highlighting the importance of testing multiple species with known differences in tolerance. We hypothesized that acidification would compound the effects of hypoxia, increasing susceptibility across all species, especially those already known to be hypoxia-sensitive.

Instead of hours-long incubations where corals naturally draw down DO to reach anoxia, we performed 392 short incubations spanning the full 0-100% DO range, fully crossed with either ambient (8.0) or low (7.6) pH_T_ treatments, and measured respiration across seven genotypes per species. Using these metabolic rates, we constructed hypoxia response curves from which we extracted the previously described hypoxia sensitivity (P_crit_) and oxyregulation (P_cmax_, T_pos_) metrics, with each metric providing insight into different aspects of how corals adjust their physiology as oxygen levels decline. ^19,28^. P_crit_ served as the primary measure of hypoxia sensitivity, whereas P_cmax_ and T_pos_ quantify complementary properties of oxyregulation that contextualize P_crit_. Recent debate regarding the relative utility of these metrics ^19,20,29,30^ motivated our inclusion of all three – best viewed as complementary ^19^ – to offer a more comprehensive view of hypoxia responses. Our findings reveal that acidification fundamentally alters coral hypoxia tolerance and produces markedly different outcomes among species. The acidification-driven increase in hypoxia sensitivity observed in several species suggests that coral communities may face critical thresholds sooner than expected, and may reduce the margin of safety for reef survival under the Paris Agreement targets ^17^.

## Methods

### Coral species and collection

This research was conducted under permit ARB-096-2023 issued by the Ministerio de Ambiente, República de Panamá. Four coral species (Fig. 2) were selected to reflect different families and morphological variety, and include hypothesized differences in hypoxia tolerance ^7,23,31^: *Agaricia tenuifolia* (Agariciidae, foliose/plate-like, sensitive), *Orbicella annularis* (Merulinidae, massive, intermediate), *Porites astreoides* (Poritidae, massive/encrusting, intermediate), and *Porites furcata* (Poritidae, branching, tolerant). Corals were collected on 25 October 2024 (*P. astreoides, A. tenuifolia*) and 31 October 2024 (*O. annularis, P. furcata*) from 5-7 m depth at Punta Caracol, Bocas del Toro, Panama. Seven visibly healthy colonies per species were selected at least 10 m apart from each other to maximize chances of sampling distinct genotypes ^32^.

**Figure 2.**
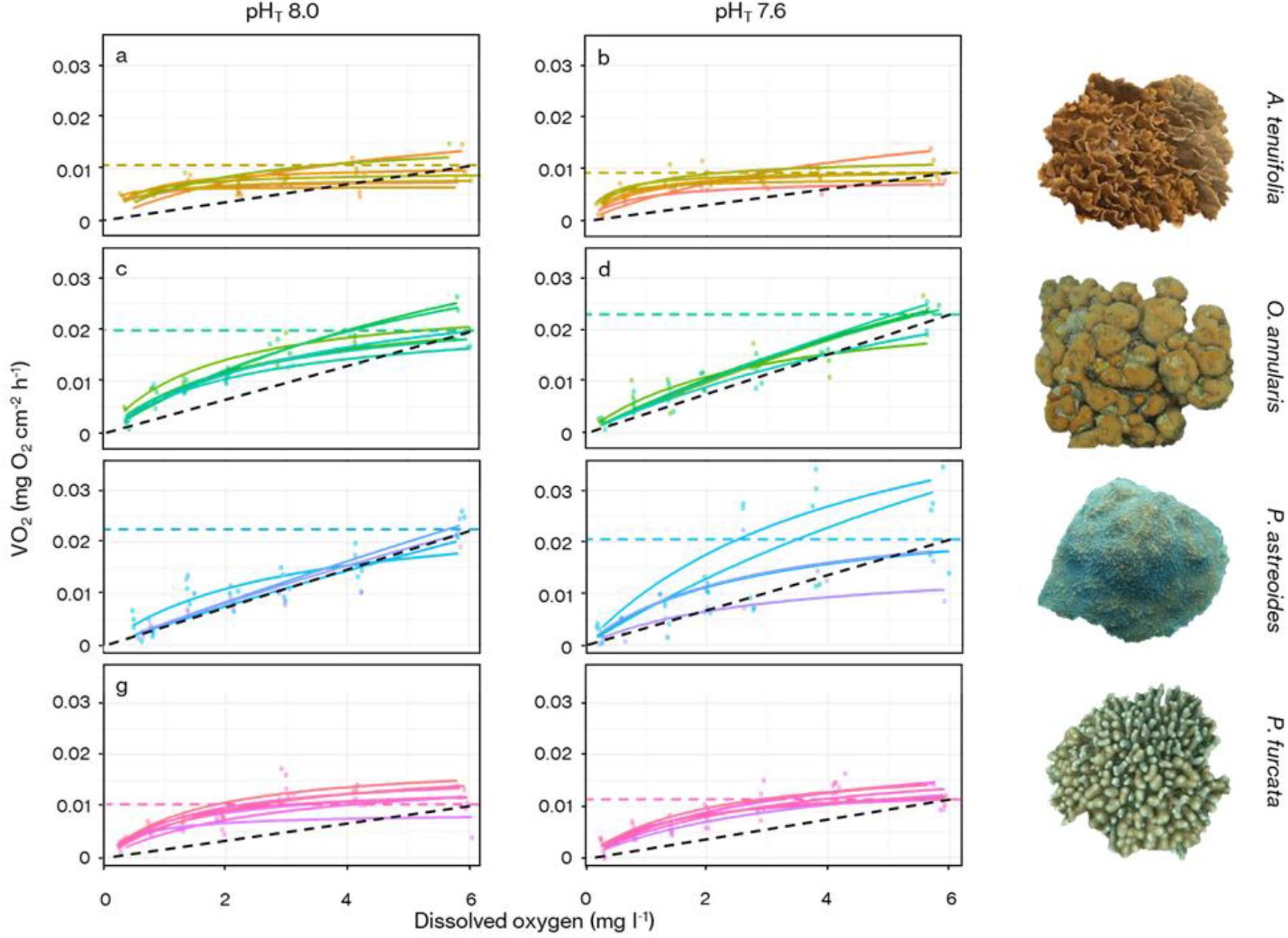
Hypoxia response curves showing inter- and intraspecific variability of the respiration rate (VO_2_) vs seawater dissolved oxygen relationship at ambient (pH_T_ 8.0; left panels) and low pH_T_ (7.6; right panels). Solid lines depict Michaelis-Menten curves fitted per genotype (n = 7 per species) for *Agaricia tenuifolia* **(a**,**b)**, *Orbicella annularis* **(c**,**d)**, *Porites astreoides* **(e**,**f)** and *Porites furcata* **(g**,**h)** measured at different dissolved oxygen concentrations. Horizontal dashed lines indicate average maximum VO_2_ (V_max_) of all measured genotypes. The black dashed line indicates perfect oxyconformity, i.e. the inability of an organism to exert any control over its metabolic rate under decreasing oxygen concentrations.

Colonies were immediately transported submerged in seawater to the Smithsonian Tropical Research Institute in Bocas del Toro in temperature-stable coolers and kept in shaded, flow-through outdoor basins (250 L) before being fragmented the following day. As collection coincided with anomalously high seawater temperatures in the area as part of the 4^th^ global bleaching event ^33^, the maximum dark-adapted quantum yield of photosystem II (Fv/Fm) was measured using a Diving PAM II fluorometer (Walz, Germany) on the night of fragmentation. These measurements confirmed that corals were not suffering chronic photo-inhibition from thermal stress (Supplementary Methods; Table S1). Colonies were cut into equally-sized fragments using a diamond band saw (Gryphon, California, USA) and hammer and chisel for a total of 14 fragments (ramets) per colony (genotype). Fragments were glued (Ecotech Coral Glue, Ecotech, Pennsylvania, USA) onto small PVC tiles. All non-coral surface area (i.e., cut edges, undersides, etc) was meticulously sealed off with glue, maintaining extreme care not to damage or cover any live coral tissue, to avoid that any live organisms other than the coral could contribute to respiration measurements. Fragments were subsequently left to recover in the outdoor basins before the start of incubations for at least 24 hours and until no signs of stress were observed (e.g., mucus production, retracted tentacles). During this period, fragments were not fed but had access to particulates between 10-50 μm that passed through the seawater system filter (Bubble Bead XF 20000 system, Propulsion Pools, MYS). Thus, metabolic rates measured here are most representative of standard metabolic rates (SMR) ^20^.

### Treatment water preparation

Seawater for incubations with the appropriate pH, temperature, and DO was prepared in an 80 L aquarium before each incubation which contained filtered seawater (Bubble Bead XF 20000 system, Propulsion Pools, MYS). pH and DO were manipulated by manually bubbling CO_2_ or N_2_ gas, respectively, under continuous mixing. Temperature and pH_T_ (Hach HQ1110 equipped with PHC101 sensor, Hach, Colorado, USA) and DO (Dissolved Oxygen Probe, Neptune Systems, California, USA) were carefully monitored until the desired composition had been achieved. pH sensors were calibrated every third day to total scale using TRIS buffer (Dickson Lab, Scripps, California, USA; ^34^.

### Metabolic oxygen incubations

Rates of metabolic oxygen consumption (i.e., dark respiration; VO_2_) were measured in full darkness on dark-adapted fragments, under various starting DO concentrations crossed with two pH levels. Starting DO concentrations were 6.07, 4.25, 3.04, 2.13, 1.52, 0.91 and 0.30 mg L^-1^ (or 100, 70, 50, 35, 25, 15 and 5% air saturation, respectively). Ambient and low pH_T_ levels were 8.0 and 7.6, respectively. These pH levels were selected to reflect pH minima during documented hypoxia events ^8^. All incubations were performed indoor, in a temperature-controlled aquarium setup (Apex A3 Pro, Neptune Systems, California, USA) programmed to maintain incubation temperature strictly at 30 °C, reflecting ambient *in situ* average seawater temperatures at Punta Caracol.

Dark-adapted single fragments of each genotype (n = 7 fragments per incubation round) were placed on raised floors in acrylic chambers (550-750 ml) filled with treatment water and fitted with magnetic stir bars. Chambers were sealed underwater to prevent the inclusion of air bubbles. Chambers were then transferred to a water bath maintaining constant water temperature, on top of magnetic stir plates to ensure constant water circulation and mixing inside the chambers. Oxygen and temperature sensors (PreSens Oxy-4 ST with PSt3 and PT100 DO and T sensors, respectively; PreSens, Regensburg, Germany) were fitted through the lids, and oxygen consumption was logged every 5 seconds across 30 minutes. Incubations at the different starting DO and pH levels were done in randomized order, across a maximum of 4 days per species (7 incubation rounds per day). Pivotally, each coral fragment was only incubated once to avoid any carry-over effects confounding respiration measurements. VO_2_ was calculated across the last 10 minutes of the 30-minute incubation to prevent potential coral handling stress influencing measurements and ensure sensor and slope stability. Blank incubations containing only treatment water were done simultaneously and used to correct VO_2_ for rates of background respiration. Post-incubation pH_T_ measurements in coral incubations showed maximum 0.02 units decrease (-0.007 ± 0.011; mean ± SD, n = 43 incubations tested) compared to the incubation start. A set of separate, trial continuous oxygen drawdown incubations for *A. tenuifolia* (n = 2), *O. annularis* (n = 3) and *P. astreoides* (n = 3; Supplementary Methods, Fig. S2) were performed using an identical setup, but were left to run until DO reached <2.5% air saturation. Pre and post-incubation pH measurements revealed that pH_T_ decreased by 0.52-0.70 units following full O_2_ drawdown.

### Coral surface area

After incubations, fragment surface area was determined using 3D photogrammetry ^35^ to provide highly accurate area measurements compared to more traditional methods. In brief, fragments with size reference were photographed from all sides at 0°, 45° and 90° angles (40-50 photos total) and 3D models were stitched using Autodesk Recap Photo software (Autodesk, California, USA). After calibration, completed models were trimmed to exclusively represent live coral tissue, and surface area was extracted and used to standardize VO_2_.

### Data and statistical analyses

Four different metrics characterizing either hypoxia sensitivity or oxyregulation capacity were calculated: (1) the critical hypoxia threshold (P_crit_) – defined as the lowest oxygen concentration at which VO_2_ remains constant ^19,36^; Fig. 1a) – represents a measure of hypoxia sensitivity as organisms cannot maintain stable respiration rates below this threshold. (2) The maximum positive oxyregulation threshold (P_cmax_) – defined as the oxygen concentration where the regulation function ρ(O_2_) reaches its highest value ^19^ – represents the oxygen concentration where the coral exerts maximum regulation to sustain respiration rates (Fig. 1b). (3) The total amount of positive oxyregulation (T_pos_) – defined as the integral of the regulation function ρ(O_2_) ^19^ – represents the total capacity of the coral to regulate respiration across the entire DO range (Fig. 1b). (4) The maximum rate of organismal respiration (V_max_), which converges with SMR if reached at normoxia ^37^. V_max_ was included to assess whether experimental treatments altered maximal metabolic capacity, ensuring that changes in hypoxia metrics were not driven by treatment-induced metabolic depression.

Replicate fragments of all genotypes were included in every incubation at each DO and pH combination. This approach allowed all metrics of oxyregulation and hypoxia sensitivity to be calculated at genotype level (n = 7 per species) in a fully balanced design, and permitted analysis of genotypic variation in hypoxia tolerance. All analyses were carried out in R (v. 4.3.3; ^38^) using RStudio (v. 2023.12.1 Build 402; RStudio Team 2024). Hypoxia response curves were created for each genotype of all four species by fitting Michaelis-Menten curves to the VO_2_ vs DO data ^28,29^. Michaelis-Menten curves were selected based on best-fit solution by AIC analysis, comparing Michaelis-Menten, hyperbolic tangent, logarithmic and linear relationships (Table S2). For 5 genotypes (including genotypes of *P. astreoides* at ambient (2) and low (2) pH, and *O. annularis* (1) under low pH), linear functions were fitted because the data exhibited non-asymptotic behavior, preventing the application of Michaelis-Menten functions. Maximum respiration rates (V_max_) were determined from the fitted curves at or near maximum DO values.

The critical oxygen threshold (P_crit_) was then calculated by applying the segmented regression analysis using the *respR* function ^39^. Regulatory functions *ρ*(O_2_) (Fig. S1) were established for each genotype from the Michaelis-Menten or linear curves fitted to the VO_2_-DO data using Equation 1 and methods described in Cobbs & Alexander (2018), where f(DO) and f’(DO) are the fitted Michaelis-Menten function and its derivative, re spectively:

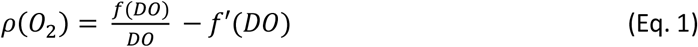

The total amount of oxyregulation (T_pos_), as well as the maximum positive oxyregulatory threshold (P_cmax_) were subsequently extracted from the regulatory function for each genotype at ambient and low pH. Differences in V_max_, P_cmax_, T_pos_, and P_crit_ thresholds between species and pH were analyzed by performing Yuen-Welch robust ANOVA with 20% trim using the WRS2 package ^40,41^, since assumptions for homogeneity and normality were violated (Table S3). Post-hoc pairwise comparisons for main effects were conducted using the lincon function ^42^, while interaction effects were tested using the mcppb20 function, applying a percentile bootstrap approach with ^43^ step-up procedure for p-value adjustment.

## Results

### Metabolic performance

Seawater pH changes were minimal during incubations (maximum –0.02 units; –0.007 ± 0.011, mean ± SD), remaining well within the range acceptable for comparisons across treatments differing by 0.4 units. Model diagnostics for the hypoxia response curves (Table S2) indicate that 7 DO starting concentrations were appropriate to fit the curves and support robust conclusions. This validates the use of short incubations at defined starting DO and pH levels for constructing hypoxia response curves. Respiration rates (VO_2_) of all coral species decreased with declining seawater DO concentration (Fig. 2), albeit differing in the rate of VO_2_ decline as well as maximum respiration (V_max_) rates. V_max_ was achieved at the highest DO (∼6 mg L^-1^ or 100% air saturation).

### Hypoxia tolerance

Acidification altered hypoxia sensitivity, as measured via P_crit_, in a species-specific manner (interactive effect between species and pH (F = 51.716, p = 0.001), Fig. 3a, Table S3). *Agaricia tenuifolia* became more tolerant to hypoxia under acidification as indicated by a 3.5 times lower P_crit_, whereas both *P. astreoides* and *P. furcata* became more sensitive (31 and 20% higher P_crit_, respectively). Only *O. annularis* maintained hypoxia tolerance independent of pH. The oxygen concentration where corals attained maximum positive oxyregulation, P_cmax_ (Fig. 3b), was unaffected by pH, but differed between species (F = 43.690, p = 0.001). *Agaricia tenuifolia* was able to sustain maximum positive oxyregulation at significantly lower DO concentration (13-68% lower) than all other species, and *P. furcata* achieved this at 51% lower DO compared to *O. annularis*. However, although *A. tenuifolia* was able to achieve positive oxyregulation at much lower DO than any other species, it still had similar total oxyregulation (T_pos_; Fig. 3c) compared to *O. annularis* and *P. furcata* owing to a flatter regulation profile (Fig. 2). Exposure to acidification negatively influenced T_pos_ in a species-specific manner (F = 25.0937, p = 0.002), reducing T_pos_ in both *O. annularis* and *P. furcata* by 64 and 23%, respectively, while remaining unchanged in the other two species. Overall, V_max_ was insensitive to changes in pH in any of the species, but differed significantly between species (Fig. 3d), being approximately 50% lower in *P. furcata* and *A. tenuifolia* compared to *P. astreoides* and *O. annularis* (F = 101.4823, p = 0.001).

**Figure 3.**
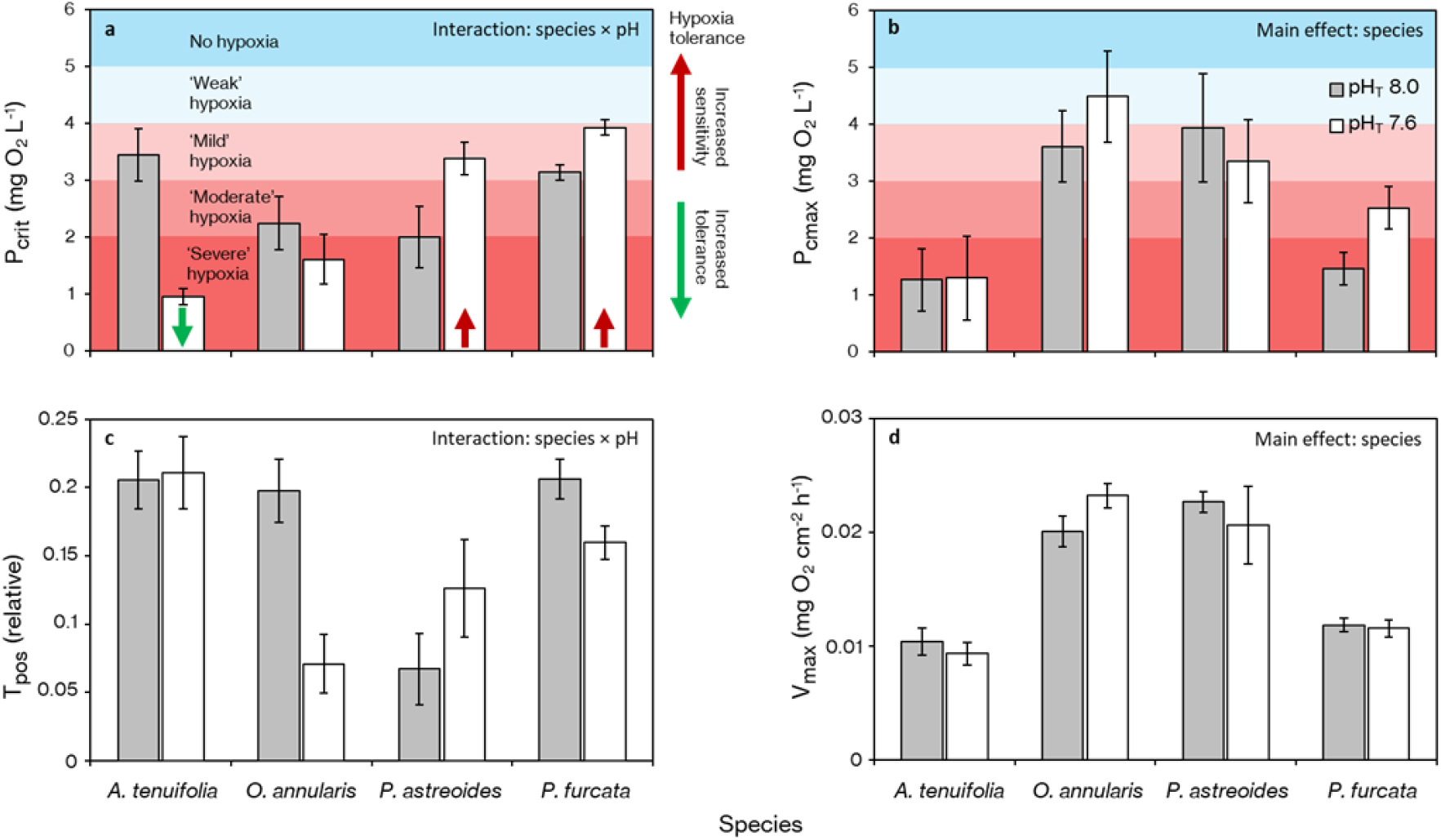
Metrics of hypoxia sensitivity and oxyregulation at ambient (pH_T_ 8.0; grey bars) and low pH_T_ (7.6; white bars). Critical hypoxia threshold (P_crit_; **a)**, DO concentration where maximum positive oxyregulation (P_cmax_; **b**) was achieved, total positive oxyregulation (T_pos_; **c)**, and maximum respiration rates (V_max_; **d)** for *Agaricia tenuifolia, Orbicella annularis, Porites astreoides*, and *Porites furcata*. V_max_ were measured directly in incubation experiments under ∼100% air saturation. P_crit_ values were determined from the inflection point of the VO_2_-DO hypoxia response curve using segmented regression analysis, whereas P_cmax_ and T_pos_ were obtained from the oxygen regulation ρ(O_2_) function derived from the hypoxia response curves. Shown is mean ± SE (n = 7 per species). Panel background colors represent deoxygenation levels identified by ^12^: No hypoxia (>5 mg O_2_ L^-1^), ‘weak’ hypoxia (4-5 mg O_2_ L^-1^), ‘mild’ hypoxia (3-4 mg O_2_ L^-1^), ‘moderate’ hypoxia (2-3 mg O_2_ L^-1^) and ‘severe’ hypoxia (<2 mg O_2_ L^-1^). Arrows inside the low-pH bars indicate significantly increased tolerance (green) or sensitivity (red), respectively, under acidification.

## Discussion

Despite the increasing prevalence of deoxygenation in coastal marine ecosystems and growing evidence that it contributes to the accelerated deterioration of coral reefs ^3,4,7^, knowledge of coral hypoxia tolerance remains limited, and largely non-existent in the context of co-occurring acidification ^1,10,22,44^. While acidification has been shown to influence hypoxia sensitivity in other marine taxa ^10,18,45^, its impact on hypoxia sensitivity of reef-building corals has remained unexplored until now. Our results indicate that acidification profoundly alters coral hypoxia tolerance and produces divergent outcomes across species. Interestingly, acidification enhanced hypoxia tolerance in *A. tenuifolia* but reduced it, to varying degrees, in the three other species. On reef scales, where pH is much more variable than in the open ocean and can be affected by coastal acidification and localized hypoxia events ^46^, this may pose a significant threat to sensitive species and drive dominance shifts in coral communities depending on species’ hypoxia and acidification tolerance. Globally, acidification may accelerate the transgression of critical hypoxia thresholds and compound the effects of warming, further undermining reef resilience ^10,17^.

The critical hypoxia threshold P_crit_ is the most widely used metric to determine hypoxia sensitivity in aquatic organisms ^20,47^, yet its application in symbiotic corals is still limited ^21–24,48^. Our P_crit_ analysis indicates that acidification can increase coral hypoxia sensitivity (Fig. 3a), i.e., it shifts the critical oxygen threshold below which metabolism cannot be maintained upwards for some coral species towards the mildest hypoxia category (classified as ‘weak hypoxia’; 4-5 mg O_2_ L^-1^) predicted for reefs worldwide under climate change ^12^. We note that this finding is not an artefact of pH-induced metabolic depression ^49^ since maximum respiration rates (V_max_) remained unaffected by acidification in all four species (Fig. 2). This is consistent with reports of unchanged coral respiration under acidification ^50^, and confirms that the observed changes in P_crit_ and oxyregulation metrics are caused by the combination of acidification and hypoxia to produce changes in hypoxia sensitivity at lower DO.

Under ambient pH, all species exhibited P_crit_ values within the ‘severe’ (<2 mg O_2_ L^-1^) to ‘moderate’ (2-3 mg O_2_ L^-1^) hypoxia categories, experienced daily by 13-34% of reefs currently ^12^. Acidification raised this threshold in *P. astreoides* and *P. furcata* into the ‘mild’ (∼3-4 mg O_2_ L^-1^) hypoxia category, representing oxygen levels already experienced daily by 50% of reefs today and by an estimated 72% of reefs under the high-emission SSP5-8.5 scenario by 2100 ^12^. These categorical hypoxia thresholds, however, are heuristic: the commonly used 2 mg O2 L^−1^ hypoxia threshold derives from temperate and pelagic systems and does not reflect the more oxygen-sensitive physiology of many marine taxa ^9,51^. Even ‘mild’ hypoxia as defined by ^12^ in their framework may therefore impose physiological stress ^52^.

While the pH levels applied here are close to, though slightly lower than those projected for the open ocean under the SSP5-8.5 scenario by the end of this century, comparable or even lower pH values already arise locally at night or during hypoxia events on present-day reefs ^8,13^. This underscores that our experimental acidification conditions are ecologically realistic under current extremes.

Moreover, while ^12^ accounted for warming-driven seawater oxygen loss, their projections did not include the direct effects of heat on hypoxia tolerance – known to further elevate P_crit_ and constrain metabolism ^1,7,53^. Together, these findings suggest that concurrent acidification and deoxygenation, alongside warming, may increasingly push corals beyond their viable oxygen envelope. Contrary to our expectations, acidification enabled *A. tenuifolia*, perceived to be a moderately hypoxia-sensitive species compared to *P. astreoides* and *P. furcata* ^7,31^, to maintain stable metabolism at much lower DO concentrations than before, almost as low as reported minimum extremes during acute hypoxia events ^3,54^. Thus, unexpected resistance to co-occurring stressors in some species complicates projections of future community composition and emphasizes the need for further multi-stressor research.

Reef-building corals are generally considered oxyregulators despite their anatomical simplicity ^22^. Here we show that acidification also influenced the capacity for metabolic oxyregulation across all species examined, although the nature and magnitude of these effects varied. Reductions in total oxyregulatory capacity (T_pos_) in *P. furcata* and *O. annularis* indicate a diminished ability to maintain stable metabolic rates under acidification (Fig. 3c). *P. furcata* also exhibited a higher oxygen threshold for maximum oxyregulation (P_cmax_), suggesting greater vulnerability to hypoxia under acidified conditions; however, because this pattern was not statistically supported in the interaction model, we interpret it with caution (Fig. 3b). In contrast, *Agaricia tenuifolia* maintained oxyregulatory performance despite acidification, though its P_crit_ was affected. These species-specific responses contrast with the generally uniform negative effects of temperature-deoxygenation interactions, where warming consistently elevates P_crit_ and suppresses oxyregulation ^1^, highlighting the complexity of acidification-driven physiological outcomes.

When comparing responses under ambient and low pH for each genotype, most individuals retained at least some oxyregulatory, with only a small subset (n=5 out of 56) exhibiting complete oxyconformity (T_pos_ ≈ 0). This variation reflects intrinsic differences in metabolic flexibility both within and among species, consistent with previous findings that, while most corals function as oxyregulators under ambient pH, the strength of this capacity varies markedly between species ^21,22^. Collectively, our comprehensive approach using multiple widely-used metrics demonstrates that acidification impacted hypoxia tolerance and/or oxyregulation in all species to some extent. This pervasive response underscores the widespread impact of acidification on coral metabolic resilience and highlights species-specific pathways through which acidification can compromise coral tolerance to low-oxygen stress. Together, these results suggest that compound deoxygenation–acidification events, though rarely considered, may drive coral community composition changes comparable to those of the more widely studied deoxygenation–warming interactions ^1,7^.

Our findings also have important methodological implications – not just for corals but aquatic organisms in general. During the continuous draw-down incubations conducted to compare pH changes between short and long incubations, pH_T_ decreased dramatically before reaching the incubation end-point (0.52-0.70 units). Because we demonstrate that hypoxia sensitivity and regulatory metrics are themselves influenced by pH, hypoxia response curves obtained from hours-long, continuous drawdown incubations are inherently confounded by low pH and do not solely capture responses to oxygen availability. Although combined pH-DO effects have been documented in other marine taxa ^10^, coral studies published to date did not include pH control nor did they report post-incubation pH measurements ^21–25,48^. Given that acidification either increased or decreased hypoxia tolerance depending on species, the likely confounding influence of unmeasured pH changes on past results remains difficult to resolve, underscoring the necessity of incorporating pH control and monitoring into future respirometry work.

Projecting coral reef trajectories under accelerating climate change remains challenging given the co-occurrence of thermal stress, deoxygenation, and ocean acidification. Our results demonstrate that acidification modulates the oxygen thresholds and oxyregulatory performance that define coral hypoxia tolerance, with responses that are strongly species-specific. The observed divergent outcomes point to the likelihood of ecological reassembly of coral communities under co-varying stressors and increasingly frequent compound events ^7,13^, with some species gaining short-term tolerance while others may lose the capacity to regulate metabolism under combined pH and oxygen stress altogether. Such shifts, while local in their manifestation ^7^, are globally significant. The acidification levels tested here, while only intermittently reached on most present-day reefs, are projected to occur with increasing frequency and duration on future reefs as ocean carbon uptake intensifies ^55^.

While compound deoxygenation–warming events are increasingly recognized as important drivers of coral mortality and ecosystem collapse ^3,13,56^, our findings indicate that deoxygenation-acidification interactions, though underrecognized, could impose comparably severe constraints on coral. By elevating coral hypoxia thresholds into oxygen concentrations already experienced daily on many reefs, ^12^, acidification may amplify the frequency and intensity of metabolic instability and push coral assemblages closer to ecological tipping points ^6,57^. These findings underline that future assessments of coral vulnerability must account for multiple, interacting climate change stressors – deoxygenation, acidification, and warming – rather than treating them in isolation, and that conservation strategies will need to incorporate species-specific physiological limits when prioritizing coral lineages and reef habitats most capable of sustaining ecosystem function under compound stress.

## Acknowledgements

We thank Rachel Collin, Plinio Gondola, and the other staff at the Smithsonian Tropical Research Institute at Bocas del Toro Research Station for their invaluable support and help, including access to the wet lab facility. We also thank Jasper de Goeij for lending oxygen and temperature sensors for the incubations, and Ben Martin for feedback on data analysis. This project was funded by a VIDI Grant from the Dutch Research Council (NWO) (to VS, grant # VI.Vidi.203.069).

## Author contributions

RZ and VS designed the study, with input from KJ. RZ, SL and KJ conducted the fieldwork, logger deployment, coral husbandry and experimental incubations. RZ analyzed the data, with contributions from VS and KJ. RZ wrote the original manuscript draft, and all authors contributed to its final form.

## Supplementary Information

### Supplementary Methods

#### Photochemical efficiency (Fv/Fm) measurements

After fragmentation of the visibly healthy coral colonies for use in the metabolic oxygen measurements and covering of all non-coral tissue fragment surface, the maximum quantum yield of photosystem II (Fv/Fm) was assessed using Pulse Amplitude Modulation (PAM) fluorometry (Diving PAM II, Walz, Germany). The maximum quantum yield of photosystem II serves as an indicator of photophysiological performance and an empirical proxy for thermal stress (i.e., bleaching). These measurements were conducted to confirm that the corals were not experiencing overt thermal stress due to the warmer than normal, ambient seawater temperatures. While these data show that corals were not experiencing chronic photoinhibition (i.e., heat stress) (Table S1), this assessment does not preclude the possibility that the coral host or its symbiotic algae experienced other sublethal or transient effects due to the elevated temperatures. Fv/Fm measurements were performed on the same day as the fragmentation, approximately 1.5 h after sunset to ensure that fragments were dark-adapted. Measurements were performed on a subset of the fragments, randomly selecting duplicate fragments from each colony (= genotype) from among the 14 fragments per genotype per species. PAM fluorometer settings were as selected as follows: Gain 1; Dampening 2; ETR-Factor 0.84; F-Offset 80; Measuring Light Intensity 11; Measuring Light Frequency 3; F0’-Mode disabled; Actinic Light Intensity 1; Actinic Light Factor 1.40; SAT-Pulse Intensity 4; SAT-Pulse Width 0.8 s.

#### Hypoxia response curves – continuous drawdown incubations

In order to determine pH changes during long, continuous, sealed incubations, hypoxia response curves ^1^ were performed with fragments of *Agaricia tenuifolia* (n = 2), *Orbicella annularis* (n = 3), and *Porites astreoides* (n = 3) under ambient starting pH_T_ and complete darkness (Fig. S2). Dissolved oxygen (DO) consumption was measured on incubated fragments from different genotypes using a PreSens Oxy-4 ST module equipped with PSt3 DO sensors (PreSens, Regensburg, Germany). During these incubations, temperature was maintained at 30.5 °C and O_2_ concentration was measured in a single continuous drawdown every 2 or 5 minutes until chamber DO reached < 2% air saturation. Incubations lasted between 3:10 h and 13:30 h depending on coral biomass and chamber volume (ranging between 540 and 767 ml). Incubations containing only seawater for estimating background respiration rates were ran parallel to the coral incubations as blanks. Starting incubation pH_T_ ranged between 8.04 and 8.06, and end-point pH_T_ values were 0.52 to 0.70 units lower (pH_T_ 7.34-7.53) in chambers containing coral, while blank incubation pH_T_ was unchanged.

Rates of oxygen consumption (VO_2_; mg O_2_ min^-1^) were calculated at 5-minute intervals from the hypoxia response curves ^2^. Michaelis-Menten type functions were fitted through the VO_2_-DO data (Fig. S2 a-c). Oxygen regulation functions ρ(O_2_) were then calculated from the hypoxia response curves ^1^ using Equation S1, where f(DO) and f’(DO) are the fitted Michaelis-Menten function and its derivative respectively (Fig. 2 d-f).

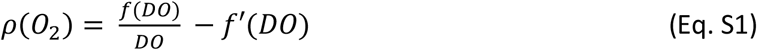

## Supplementary Tables

**Table S1.**
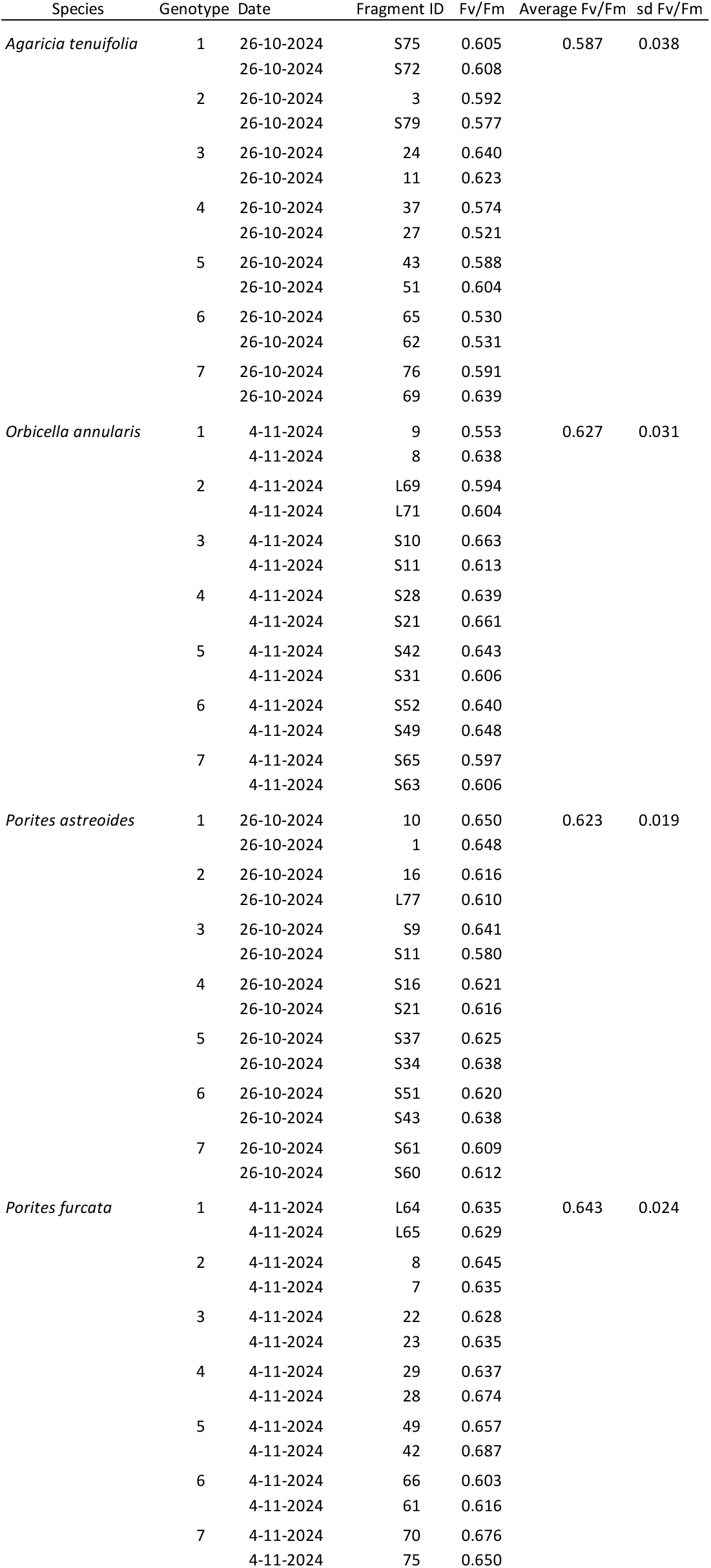
Maximum quantum yield of photosystem II (Fv/Fm) measured on duplicate dark-adapted, randomly selected fragments of each genotype (n = 7 genotypes) of the four experimental coral species. Fv/Fm was measured after fragmentation and covering non-coral surface areas, but before respirometry incubations.

| Species | Genotype | Date | Fragment ID | Fv/Fm | Average Fv/Fm | sd Fv/Fm |
| --- | --- | --- | --- | --- | --- | --- |
| *Agaricia tenuifolia* | 1 | 26-10-2024 | S75 | 0.605 | 0.587 | 0.038 |
|  |  | 26-10-2024 | S72 | 0.608 |  |  |
|  | 2 | 26-10-2024 | 3 | 0.592 |  |  |
|  |  | 26-10-2024 | S79 | 0.577 |  |  |
|  | 3 | 26-10-2024 | 24 | 0.640 |  |  |
|  |  | 26-10-2024 | 11 | 0.623 |  |  |
|  | 4 | 26-10-2024 | 37 | 0.574 |  |  |
|  |  | 26-10-2024 | 27 | 0.521 |  |  |
|  | 5 | 26-10-2024 | 43 | 0.588 |  |  |
|  |  | 26-10-2024 | 51 | 0.604 |  |  |
|  | 6 | 26-10-2024 | 65 | 0.530 |  |  |
|  |  | 26-10-2024 | 62 | 0.531 |  |  |
|  | 7 | 26-10-2024 | 76 | 0.591 |  |  |
|  |  | 26-10-2024 | 69 | 0.639 |  |  |
| *Orbicella annularis* | 1 | 4-11-2024 | 9 | 0.553 | 0.627 | 0.031 |
|  |  | 4-11-2024 | 8 | 0.638 |  |  |
|  | 2 | 4-11-2024 | L69 | 0.594 |  |  |
|  |  | 4-11-2024 | L71 | 0.604 |  |  |
|  | 3 | 4-11-2024 | S10 | 0.663 |  |  |
|  |  | 4-11-2024 | S11 | 0.613 |  |  |
|  | 4 | 4-11-2024 | S28 | 0.639 |  |  |
|  |  | 4-11-2024 | S21 | 0.661 |  |  |
|  | 5 | 4-11-2024 | S42 | 0.643 |  |  |
|  |  | 4-11-2024 | S31 | 0.606 |  |  |
|  | 6 | 4-11-2024 | S52 | 0.640 |  |  |
|  |  | 4-11-2024 | S49 | 0.648 |  |  |
|  | 7 | 4-11-2024 | S65 | 0.597 |  |  |
|  |  | 4-11-2024 | S63 | 0.606 |  |  |
| *Porites astreoides* | 1 | 26-10-2024 | 10 | 0.650 | 0.623 | 0.019 |
|  |  | 26-10-2024 | 1 | 0.648 |  |  |
|  | 2 | 26-10-2024 | 16 | 0.616 |  |  |
|  |  | 26-10-2024 | L77 | 0.610 |  |  |
|  | 3 | 26-10-2024 | S9 | 0.641 |  |  |
|  |  | 26-10-2024 | S11 | 0.580 |  |  |
|  | 4 | 26-10-2024 | S16 | 0.621 |  |  |
|  |  | 26-10-2024 | S21 | 0.616 |  |  |
|  | 5 | 26-10-2024 | S37 | 0.625 |  |  |
|  |  | 26-10-2024 | S34 | 0.638 |  |  |
|  | 6 | 26-10-2024 | S51 | 0.620 |  |  |
|  |  | 26-10-2024 | S43 | 0.638 |  |  |
|  | 7 | 26-10-2024 | S61 | 0.609 |  |  |
|  |  | 26-10-2024 | S60 | 0.612 |  |  |
| *Porites furcata* | 1 | 4-11-2024 | L64 | 0.635 | 0.643 | 0.024 |
|  |  | 4-11-2024 | L65 | 0.629 |  |  |
|  | 2 | 4-11-2024 | 8 | 0.645 |  |  |
|  |  | 4-11-2024 | 7 | 0.635 |  |  |
|  | 3 | 4-11-2024 | 22 | 0.628 |  |  |
|  |  | 4-11-2024 | 23 | 0.635 |  |  |
|  | 4 | 4-11-2024 | 29 | 0.637 |  |  |
|  |  | 4-11-2024 | 28 | 0.674 |  |  |
|  | 5 | 4-11-2024 | 49 | 0.657 |  |  |
|  |  | 4-11-2024 | 42 | 0.687 |  |  |
|  | 6 | 4-11-2024 | 66 | 0.603 |  |  |
|  |  | 4-11-2024 | 61 | 0.616 |  |  |
|  | 7 | 4-11-2024 | 70 | 0.676 |  |  |
|  |  | 4-11-2024 | 75 | 0.650 |  |  |

**Table S2.** Model comparison for best-fit curves depicting the relationship between respiration rate (VO_2_) and dissolved oxygen concentration in *Agaricia tenuifolia, Orbicella annularis, Porites astreoides* and *Porites furcata* across ambient (8.0) and low (7.6) pH_T_. Comparisons included Michaelis-Menten (MM), hyperbolic tangent (Tanh), logarithmic (Log) and linear relationships. Curve fitting occurred on combined VO_2_ vs oxygen data per species per pH, and ranked from best to worst. In three cases the relationship was best described by a linear response, whereas in all other cases a non-linear fit was most appropriate. Ultimately, Michaelis-Menten relationships were selected to depict the relationship in all cases because these never ranked worse than second place.

| Species | $\text{pH}_T$ | Best-fit | | | 2<sup>nd</sup> best-fit | | | 3<sup>rd</sup> best-fit | | | 4<sup>th</sup> best-fit | | |
| --- | --- | --- | --- | --- | --- | --- | --- | --- | --- | --- | --- | --- | --- |
| | | Model | $R^2$ | AIC | Model | $R^2$ | AIC | Model | $R^2$ | AIC | Model | $R^2$ | AIC |
| *A. tenuifolia* | 8.0 | Linear | 0.493 | -477.094 | Log | 0.477 | -475.620 | MM | 0.396 | -468.515 | Tanh | 0.265 | -458.949 |
|  | 7.6 | Tanh | 0.630 | -476.285 | MM | 0.630 | -476.242 | Log | 0.551 | -466.791 | Linear | 0.435 | -455.577 |
| *O. annularis* | 8.0 | MM | 0.869 | -454.790 | Log | 0.867 | -454.078 | Tanh | 0.862 | -452.268 | Linear | 0.803 | -434.960 |
|  | 7.6 | Linear | 0.879 | -447.398 | MM | 0.867 | -442.884 | Tanh | 0.860 | -440.265 | Log | 0.839 | -433.555 |
| *P. astreoides* | 8.0 | Linear | 0.734 | -407.620 | MM | 0.721 | -405.359 | Tanh | 0.721 | -405.351 | Log | 0.662 | -395.885 |
|  | 7.6 | Tanh | 0.547 | -351.122 | MM | 0.537 | -350.102 | Log | 0.533 | -349.722 | Linear | 0.505 | -346.856 |
| *P. furcata* | 8.0 | Tanh | 0.677 | -453.041 | MM | 0.657 | -450.081 | Log | 0.612 | -444.069 | Linear | 0.482 | -429.870 |
|  | 7.6 | Tanh | 0.779 | -465.103 | MM | 0.763 | -461.663 | Log | 0.750 | -458.988 | Linear | 0.654 | -443.112 |

**Table S3.**
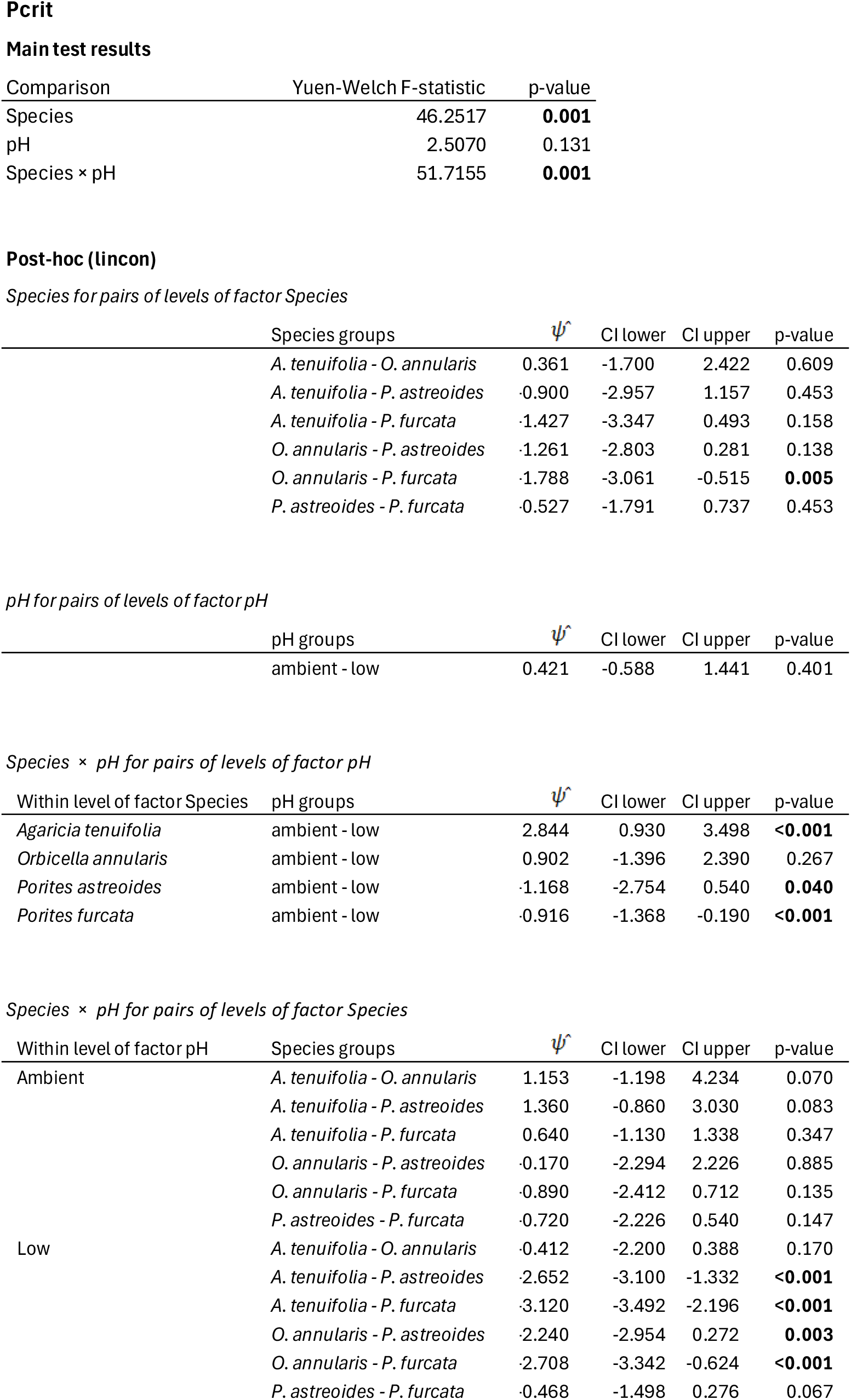
Yuen-Welch type ANOVA results for the critical hypoxia threshold (P_crit_), the DO threshold below which stable metabolic rates can no longer be maintained, and post-hoc analyses indicating differences between groups. Analyses were done using standard 20% trimmed means and Winsorized variances. In the post-hoc comparisons, values for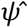(Psi-hat) indicate absolute differences between model-estimated group means. Significant outcomes are indicated in bold typeface.

Main test results
| Comparison | Yuen-Welch F-statistic | p-value |
| --- | --- | --- |
| Species | 46.2517 | **0.001** |
| pH | 2.5070 | 0.131 |
| Species × pH | 51.7155 | **0.001** |

Post-hoc (lincon) Species for pairs of levels of factor Species
| Species groups | $\psi^{\wedge}$ | CI lower | CI upper | p-value |
| --- | --- | --- | --- | --- |
| *A. tenuifolia* - *O. annularis* | 0.361 | -1.700 | 2.422 | 0.609 |
| *A. tenuifolia* - *P. astreoides* | -0.900 | -2.957 | 1.157 | 0.453 |
| *A. tenuifolia* - *P. furcata* | -1.427 | -3.347 | 0.493 | 0.158 |
| *O. annularis* - *P. astreoides* | -1.261 | -2.803 | 0.281 | 0.138 |
| *O. annularis* - *P. furcata* | -1.788 | -3.061 | -0.515 | **0.005** |
| *P. astreoides* - *P. furcata* | -0.527 | -1.791 | 0.737 | 0.453 |

pH for pairs of levels of factor pH
| pH groups | $\psi^{\wedge}$ | CI lower | CI upper | p-value |
| --- | --- | --- | --- | --- |
| ambient - low | 0.421 | -0.588 | 1.441 | 0.401 |

Species × pH for pairs of levels of factor pH
| Within level of factor Species | pH groups | $\psi^{\wedge}$ | CI lower | CI upper | p-value |
| --- | --- | --- | --- | --- | --- |
| *Agaricia tenuifolia* | ambient - low | 2.844 | 0.930 | 3.498 | **<0.001** |
| *Orbicella annularis* | ambient - low | 0.902 | -1.396 | 2.390 | 0.267 |
| *Porites astreoides* | ambient - low | -1.168 | -2.754 | 0.540 | **0.040** |
| *Porites furcata* | ambient - low | -0.916 | -1.368 | -0.190 | **<0.001** |

| Within level of factor pH | Species groups | $\psi^{\wedge}$ | CI lower | CI upper | p-value |
| --- | --- | --- | --- | --- | --- |
| Ambient | *A. tenuifolia* - *O. annularis* | 1.153 | -1.198 | 4.234 | 0.070 |
|  | *A. tenuifolia* - *P. astreoides* | 1.360 | -0.860 | 3.030 | 0.083 |
|  | *A. tenuifolia* - *P. furcata* | 0.640 | -1.130 | 1.338 | 0.347 |
|  | *O. annularis* - *P. astreoides* | -0.170 | -2.294 | 2.226 | 0.885 |
|  | *O. annularis* - *P. furcata* | -0.890 | -2.412 | 0.712 | 0.135 |
|  | *P. astreoides* - *P. furcata* | -0.720 | -2.226 | 0.540 | 0.147 |
| Low | *A. tenuifolia* - *O. annularis* | -0.412 | -2.200 | 0.388 | 0.170 |
|  | *A. tenuifolia* - *P. astreoides* | -2.652 | -3.100 | -1.332 | **<0.001** |
|  | *A. tenuifolia* - *P. furcata* | -3.120 | -3.492 | -2.196 | **<0.001** |
|  | *O. annularis* - *P. astreoides* | -2.240 | -2.954 | 0.272 | **0.003** |
|  | *O. annularis* - *P. furcata* | -2.708 | -3.342 | -0.624 | **<0.001** |
|  | *P. astreoides* - *P. furcata* | -0.468 | -1.498 | 0.276 | 0.067 |

**Table S4.** Yuen-Welch type ANOVA results for the DO threshold where maximum positive oxyregulation (P_cmax_) was achieved, and post-hoc analyses indicating differences between groups. Analyses were done using standard 20% trimmed means and Winsorized variances. In the post-hoc comparisons, values for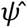 (Psi-hat) indicate absolute differences between model-estimated group means. Significant outcomes are indicated in bold typeface.

Main test results
| Comparison | Yuen-Welch F-statistic | p-value |
| --- | --- | --- |
| Species | 43.6900 | **0.001** |
| pH | 0.6777 | 0.423 |
| Species × pH | 5.6069 | 0.208 |

Post-hoc (lincon) Species for pairs of levels of factor Species
| Species groups | $\psi^{\wedge}$ | CI lower | CI upper | p-value |
| --- | --- | --- | --- | --- |
| *A. tenuifolia* - *O. annularis* | -3.517 | -5.635 | -1.399 | **0.001** |
| *A. tenuifolia* - *P. astreoides* | -3.097 | -5.747 | -0.447 | **0.018** |
| *A. tenuifolia* - *P. furcata* | -1.176 | -2.346 | -0.007 | **0.026** |
| *O. annularis* - *P. astreoides* | 0.420 | -2.581 | 3.422 | 0.685 |
| *O. annularis* - *P. furcata* | 2.341 | 0.213 | 4.469 | **0.021** |
| *P. astreoides* - *P. furcata* | 1.921 | -0.735 | 4.577 | 0.089 |

pH for pairs of levels of factor pH
| pH groups | $\psi^{\wedge}$ | CI lower | CI upper | p-value |
| --- | --- | --- | --- | --- |
| ambient - low | -0.587 | -2.285 | 1.111 | 0.487 |

Species × pH for pairs of levels of factor pH
| Within level of factor Species | pH groups | $\psi^{\wedge}$ | CI lower | CI upper | p-value |
| --- | --- | --- | --- | --- | --- |
| *Agaricia tenuifolia* | ambient - low | 0.246 | -2.314 | 2.400 | 0.881 |
| *Orbicella annularis* | ambient - low | -1.564 | -3.506 | 2.365 | 0.264 |
| *Porites astreoides* | ambient - low | 0.732 | -3.844 | 4.154 | 0.588 |
| *Porites furcata* | ambient - low | -1.114 | -2.538 | 0.480 | **0.027** |

| Within level of factor pH | Species groups | $\psi^{\wedge}$ | CI lower | CI upper | p-value |
| --- | --- | --- | --- | --- | --- |
| Ambient | *A. tenuifolia* - *O. annularis* | -2.612 | -5.278 | -0.041 | **<0.001** |
|  | *A. tenuifolia* - *P. astreoides* | -3.353 | -5.564 | 1.500 | **0.020** |
|  | *A. tenuifolia* - *P. furcata* | -0.438 | -2.082 | 2.140 | 0.521 |
|  | *O. annularis* - *P. astreoides* | -0.740 | -3.701 | 3.535 | 0.594 |
|  | *O. annularis* - *P. furcata* | 2.174 | 0.471 | 4.847 | **<0.001** |
|  | *P. astreoides* - *P. furcata* | 2.915 | -0.564 | 4.917 | **0.023** |
| Low | *A. tenuifolia* - *O. annularis* | -4.422 | -5.317 | -0.164 | **<0.001** |
|  | *A. tenuifolia* - *P. astreoides* | -2.866 | -4.866 | 0.510 | **0.017** |
|  | *A. tenuifolia* - *P. furcata* | -1.798 | -3.149 | 0.954 | 0.093 |
|  | *O. annularis* - *P. astreoides* | 1.556 | -2.141 | 4.913 | 0.220 |
|  | *O. annularis* - *P. furcata* | 2.625 | -0.867 | 4.044 | **0.027** |
|  | *P. astreoides* - *P. furcata* | 1.069 | -1.794 | 3.306 | 0.274 |

**Table S5.** Yuen-Welch type ANOVA results for the total positive oxyregulation (T_pos_), and post-hoc analyses indicating differences between groups. Analyses were done using standard 20% trimmed means and Winsorized variances. In the post-hoc comparisons, values for 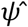 (Psi-hat) indicate absolute differences between model-estimated group means. Significant outcomes are indicated in bold typeface.

Main test results
| Comparison | Yuen-Welch F-statistic | p-value |
| --- | --- | --- |
| Species | 40.4101 | **0.001** |
| pH | 2.2694 | 0.155 |
| Species × pH | 25.0937 | **0.002** |

Post-hoc (lincon) Species for pairs of levels of factor Species
| Species groups | $\psi^{\wedge}$ | CI lower | CI upper | p-value |
| --- | --- | --- | --- | --- |
| *A. tenuifolia* - *O. annularis* | 0.084 | -0.009 | 0.178 | 0.065 |
| *A. tenuifolia* - *P. astreoides* | 0.127 | 0.026 | 0.229 | **0.012** |
| *A. tenuifolia* - *P. furcata* | 0.032 | -0.028 | 0.091 | 0.268 |
| *O. annularis* - *P. astreoides* | 0.043 | -0.074 | 0.161 | 0.297 |
| *O. annularis* - *P. furcata* | -0.053 | -0.144 | 0.039 | 0.268 |
| *P. astreoides* - *P. furcata* | -0.096 | -0.195 | 0.004 | 0.059 |

pH for pairs of levels of factor pH
| pH groups | $\psi^{\wedge}$ | CI lower | CI upper | p-value |
| --- | --- | --- | --- | --- |
| ambient - low | 0.036 | -0.015 | 0.088 | 0.163 |

Species × pH for pairs of levels of factor pH
| Within level of factor Species | pH groups | $\psi^{\wedge}$ | CI lower | CI upper | p-value |
| --- | --- | --- | --- | --- | --- |
| *Agaricia tenuifolia* | ambient - low | -0.006 | -0.124 | 0.120 | 0.855 |
| *Orbicella annularis* | ambient - low | 0.130 | 0.013 | 0.228 | **<0.001** |
| *Porites astreoides* | ambient - low | -0.081 | -0.198 | 0.071 | 0.147 |
| *Porites furcata* | ambient - low | 0.051 | -0.022 | 0.103 | **0.010** |

| Within level of factor pH | Species groups | $\psi^{\wedge}$ | CI lower | CI upper | p-value |
| --- | --- | --- | --- | --- | --- |
| Ambient | *A. tenuifolia* - *O. annularis* | 0.021 | -0.083 | 0.094 | 0.678 |
|  | *A. tenuifolia* - *P. astreoides* | 0.165 | 0.033 | 0.230 | **<0.001** |
|  | *A. tenuifolia* - *P. furcata* | -0.001 | -0.093 | 0.077 | 0.962 |
|  | *O. annularis* - *P. astreoides* | 0.144 | 0.013 | 0.232 | **<0.001** |
|  | *O. annularis* - *P. furcata* | -0.021 | -0.095 | 0.096 | 0.494 |
|  | *P. astreoides* - *P. furcata* | -0.166 | -0.215 | -0.048 | **<0.001** |
| Low | *A. tenuifolia* - *O. annularis* | 0.157 | 0.042 | 0.232 | **<0.001** |
|  | *A. tenuifolia* - *P. astreoides* | 0.090 | -0.036 | 0.227 | 0.093 |
|  | *A. tenuifolia* - *P. furcata* | 0.057 | -0.066 | 0.137 | 0.070 |
|  | *O. annularis* - *P. astreoides* | -0.067 | -0.224 | 0.090 | 0.154 |
|  | *O. annularis* - *P. furcata* | -0.100 | -0.175 | -0.006 | **<0.001** |
|  | *P. astreoides* - *P. furcata* | -0.033 | -0.170 | 0.096 | 0.487 |

**Table S6.** Yuen-Welch type ANOVA results for the maximum respiration rate V_max_, and post-hoc analyses indicating differences between groups. Analyses were done using standard 20% trimmed means and Winsorized variances. In the post-hoc comparisons, values for 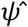(Psi-hat) indicate absolute differences between model-estimated group means. Significant outcomes are indicated in bold typeface.

Main test results
| Comparison | Yuen-Welch F-statistic | p-value |
| --- | --- | --- |
| Species | 101.4823 | **0.001** |
| pH | 0.0001 | 0.994 |
| Species × pH | 4.1143 | 0.331 |

Post-hoc (lincon) Species for pairs of levels of factor Species
| Species groups | $\psi^{\wedge}$ | CI lower | CI upper | p-value |
| --- | --- | --- | --- | --- |
| *A. tenuifolia* - *O. annularis* | -0.012 | -0.017 | -0.008 | **<0.001** |
| *A. tenuifolia* - *P. astreoides* | -0.012 | -0.018 | -0.007 | **<0.001** |
| *A. tenuifolia* - *P. furcata* | -0.002 | -0.006 | 0.001 | 0.123 |
| *O. annularis* - *P. astreoides* | -0.002 | -0.006 | 0.006 | 0.932 |
| *O. annularis* - *P. furcata* | 0.010 | 0.006 | 0.014 | **<0.001** |
| *P. astreoides* - *P. furcata* | 0.010 | 0.005 | 0.015 | **<0.001** |

pH for pairs of levels of factor pH
| pH groups | $\psi^{\wedge}$ | CI lower | CI upper | p-value |
| --- | --- | --- | --- | --- |
| ambient - low | 0.001 | -0.004 | 0.005 | 0.773 |

Species × pH for pairs of levels of factor pH
| Within level of factor Species | pH groups | $\psi^{\wedge}$ | CI lower | CI upper | p-value |
| --- | --- | --- | --- | --- | --- |
| *Agaricia tenuifolia* | ambient - low | 0.001 | -0.003 | 0.007 | 0.558 |
| *Orbicella annularis* | ambient - low | -0.004 | -0.008 | 0.005 | 0.073 |
| *Porites astreoides* | ambient - low | 0.002 | -0.012 | 0.013 | 0.611 |
| *Porites furcata* | ambient - low | 0.001 | -0.003 | 0.003 | 0.798 |

Species × pH for pairs of levels of factor Species
| Within level of factor pH | Species groups | $\psi^{\wedge}$ | CI lower | CI upper | p-value |
| --- | --- | --- | --- | --- | --- |
| Ambient | *A. tenuifolia* - *O. annularis* | -0.009 | -0.017 | 0.0004 | **<0.001** |
|  | *A. tenuifolia* - *P. astreoides* | -0.013 | -0.016 | -0.007 | **<0.001** |
|  | *A. tenuifolia* - *P. furcata* | -0.002 | -0.005 | 0.003 | 0.304 |
|  | *O. annularis* - *P. astreoides* | -0.003 | -0.008 | 0.003 | 0.104 |
|  | *O. annularis* - *P. furcata* | 0.008 | 0.003 | 0.014 | **<0.001** |
|  | *P. astreoides* - *P. furcata* | 0.011 | 0.007 | 0.015 | **<0.001** |
| Low | *A. tenuifolia* - *O. annularis* | -0.014 | -0.018 | -0.009 | **<0.001** |
|  | *A. tenuifolia* - *P. astreoides* | -0.011 | -0.024 | -0.001 | **<0.001** |
|  | *A. tenuifolia* - *P. furcata* | -0.003 | -0.006 | 0.002 | 0.100 |
|  | *O. annularis* - *P. astreoides* | 0.003 | -0.010 | 0.014 | 0.487 |
|  | *O. annularis* - *P. furcata* | 0.012 | -0.007 | 0.016 | **<0.001** |
|  | *P. astreoides* - *P. furcata* | 0.009 | -0.009 | 0.022 | **0.017** |

## Supplementary Figures

**Figure S1.**
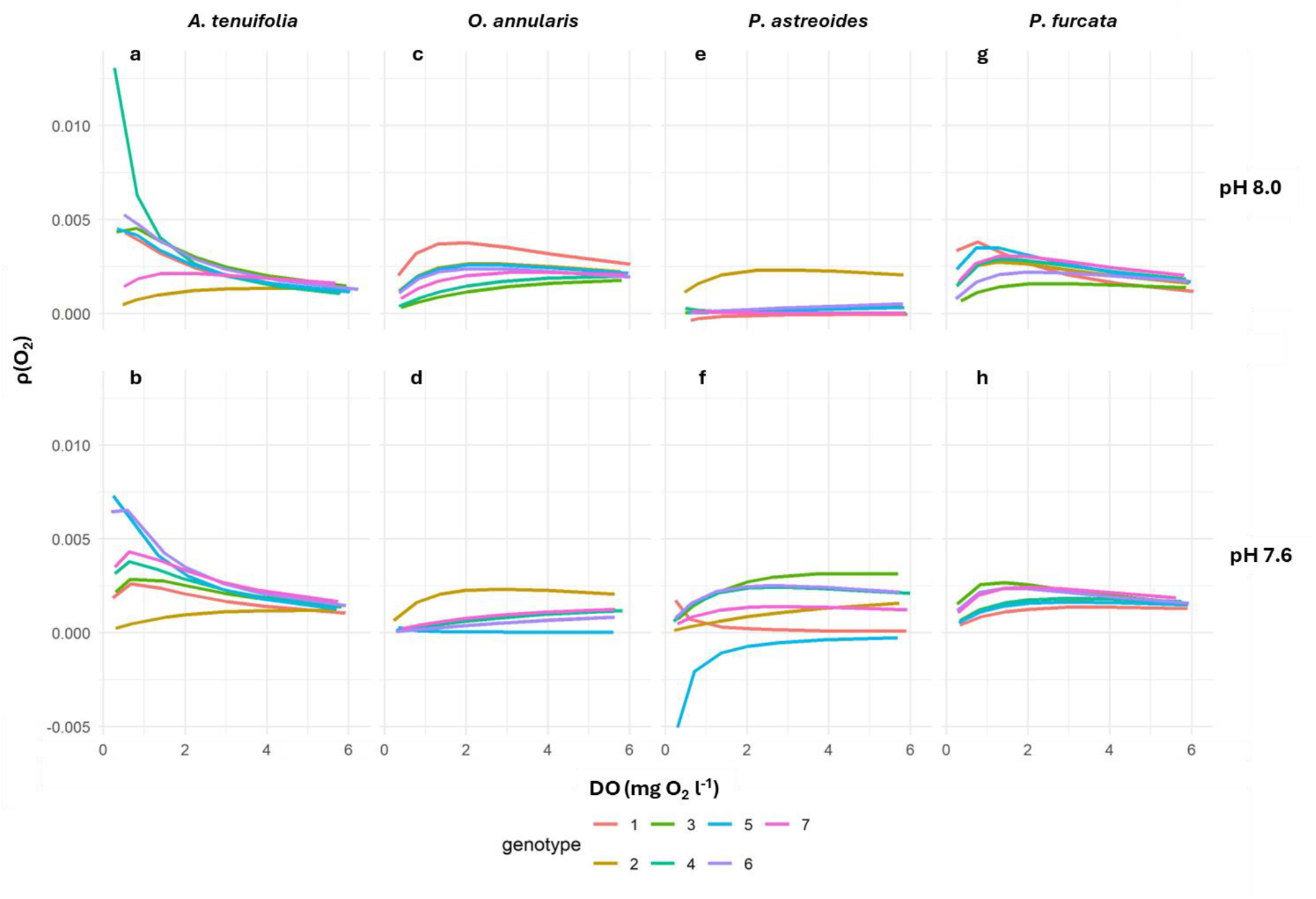
Regulation profiles illustrating the direction and strength of oxyregulation (ρ(O_2_)) across increasing *p*O_2_ for all genotypes of four coral species (*Agaricia tenuifolia, Orbicella annularis, Porites astreoides*, and *Porites furcata*), under ambient pH (pH_T_ 8.0) and acidification (pH_T_ 7.6). Regulation profiles are calculated from the Michaelis-Menten curves fitted through the VO_2_-DO data. Corresponding DO values where ρ(O_2_) reaches maximum value indicate the maximum positive oxyregulation threshold P_cmax_.

**Figure S2.**
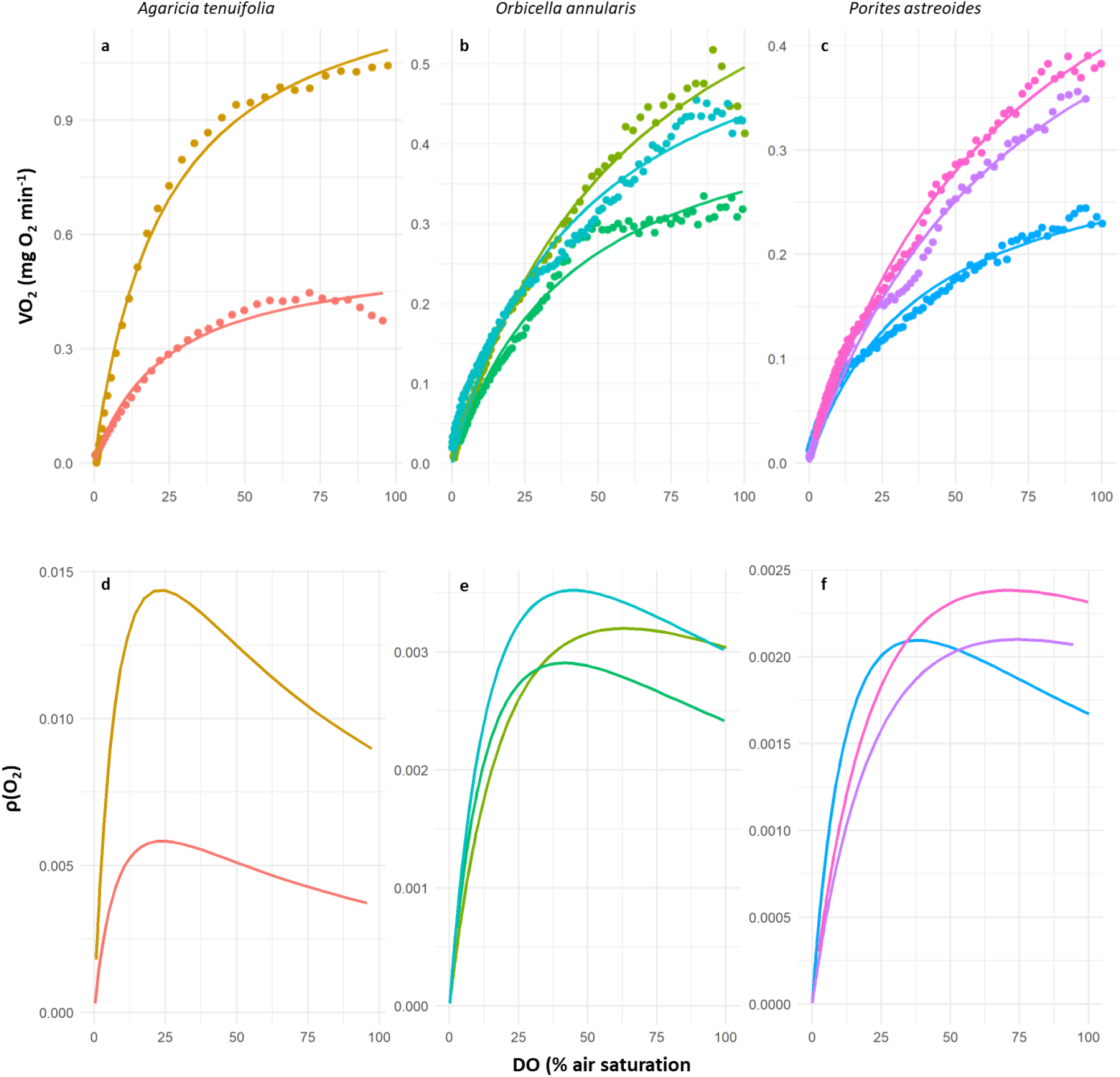
Hypoxia response curves for the long, continuous oxygen drawdown incubations (**a-c**; closed symbols) for genotypes of *Agaricia tenuifolia* (n = 2), *Orbicella annularis* (n = 3), and *Porites astreoides* (n = 3) measured at ambient temperature (30.5 °C) and pH_T_ (pH_T_ 8.04-8.06). Incubations were started at 100% oxygen saturation, and continued uninterrupted until DO concentration reached <2.5% air saturation. Metabolic oxygen rates (VO_2_) were calculated at 5-minute intervals. Michaelis-Menten type equations (solid lines) were fitted through the data to obtain oxyregulation curves. Associated oxyregulation curves ρ(O_2_) **(d-f)** corresponding to the hypoxia response curves of the same genotypes from panels a-c. No hypoxia response curves were conducted for *Porites furcata*. Note the different scales on the y-axes.

